# Microstructural meal pattern analysis reveals that nicotine is a potent anti-anorectic drug despite producing long-term anorexigenic effects

**DOI:** 10.1101/2020.02.04.934414

**Authors:** Kokila Shankar, Frederic Ambroggi, Olivier George

## Abstract

Nicotine consumption in both human and animal studies has been strongly associated with changes in feeding-related behaviors and metabolism. The current dogma is that chronic nicotine decreases food intake and increases metabolism, leading to lower body weight. However, the effect of acute nicotine intake on feeding is unclear. The present study employed microstructural and macrostructural behavioral analyses to elucidate changes in feeding behavior in animals that intravenously self-administered nicotine. At the microstructural level (seconds to minutes), nicotine increased feeding and drinking behavior during the first 5 minutes after nicotine self-administration. This effect was also observed in animals that passively received nicotine, but the effect was not observed in animals that self-administered saline or passively received saline. At the macrostructural level (hours to days), nicotine decreased body weight gain, decreased feeding, and was associated with increases in feeding and body weight gain during abstinence. These results suggest that nicotine first produces anti-anorectic effects before producing long-term anorexigenic effects. These results challenge the notion that nicotine is an anorexigenic drug and paradoxically suggest that the anorexigenic effects of nicotine may be a long-term consequence of acute anti-anorectic effects of nicotine.

## Introduction

Although cigarette smoking and tobacco use have been primarily reported to occur because of the psychoactive properties of nicotine, robust evidence suggests that tobacco is also used for its effects on body weight. Most chronic smokers exhibit lower body weight gain compared with nonsmokers. Following smoking cessation, smokers often report robust weight gain that contributes to relapse.^1–3^

Chronic exposure to nicotine or tobacco smoke in humans and nonhuman animals reduces body weight and food intake and increases metabolism, contributing to the decrease in body weight.^4–8^ The dogma for the past 30 years is that these effects are produced by the acute activation of nicotinic receptors in the brain and body.^9,10^ Preclinical models have consistently shown that nicotine decreases food intake hours and days after initiating chronic nicotine exposure; however, these studies did not report the initial (acute) effect of nicotine on feeding. Moreover, few to no studies have reported nicotine-induced decreases in food intake when given acutely in humans. One study reported an increase in caloric intake after acute nicotine exposure in humans.^11^ Nicotine reaches the brain in ∼7 seconds after a single cigarette puff,^12^ and brain nicotinic receptors are saturated after only three puffs.^13^ Furthermore, most nicotinic receptors desensitize in minutes, suggesting that if nicotine has anorectic properties, then it should produce a robust decrease in hunger or caloric intake minutes after administration. However, studies to date have not evaluated the effects of nicotine on feeding at the microstructural level (i.e., seconds to minutes after exposure to nicotine).

Microstructural meal pattern analysis is a technique that is commonly used in feeding behavior studies that can be employed to analyze acute behavioral changes.^14,15^ This approach measures behavioral changes within a small timescale, usually seconds to minutes, as a response to a specific singular event. For example, microstructural meal pattern analysis shows that the acute administration of cocaine, a potent anorectic drug, decreases food intake in rats^16^; however, such a microstructural approach has not yet been tested with nicotine.

To address this gap in the literature and identify the acute and long-term effects of nicotine on caloric intake, feeding behavior, and body weight, we performed both microstructural (seconds to minutes) and macrostructural (hours to days) analyses of rats that intravenously self-administered nicotine. Based on the current dogma, we hypothesized that acute and chronic nicotine self-administration decreases caloric intake, feeding, and body weight.

## Methods

### Animals

Male Wistar rats (250-275 g; Charles River, Hollister, CA, USA) were used for the experiments. The animals were group-housed and maintained on a 12 h/12 h light/dark cycle (lights off at 10:00 AM) with *ad libitum* access to food (45 mg grain-based tablets, TestDiet, St. Louis, MO, USA) and tap water. All of the animal procedures were approved by The Scripps Research Institute Institutional Animal Care and Use Committee and were in accordance with National Institutes of Health guidelines.

### Drugs

Nicotine hydrogen tartrate salt (Sigma, Natick, MA, USA) was dissolved in saline (pH 7.4) and self-administered via an indwelling intravenous jugular catheter. Doses are expressed as free base.

### Nicotine self-administration

The apparatus and detailed procedures for both intravenous catheterization and nicotine self-administration have been described previously.^17^ The rats were first trained to nosepoke for food and water in 23 h sessions before and after recovery from the surgical implantation of jugular catheters but were not trained to respond at the lever that was associated with nicotine delivery. Following the acquisition of these operant responses, the active and inactive levers were extended, and the rats were allowed to self-administer nicotine (0.03 mg/kg/100 μl/1 s, free base, fixed-ratio 1 [FR1], timeout [TO] 20 s) by pressing the active lever. The rats were first given access to nicotine for 1 h per day during the dark cycle (beginning at 10:00 AM) for 1 week and then separated into two groups that were given short access (ShA; 1 h/day) and long access (LgA; 23 h/day) to nicotine for 2 weeks. Another group of animals was also trained to self-administer 0.9% saline solution as a control. Passive nicotine administration was conducted using the MedPC program (MedAssociates, St. Albans, VT, USA) after habituation but before nicotine self-administration, in which one saline infusion, one nicotine infusion, or three nicotine infusions were administered to the animals in a Latin-square design.

### Peristimulus time histogram generation

Total food, water, and drug self-administration events and total active and inactive lever presses for each animal during each day of the experiment were extracted from MedPC and stored as .txt files. The files were imported into Microsoft Excel and then batch imported into Spike2 (Cambridge Electronic Design, Cambridge, United Kingdom). To generate peristimulus time histograms, the average number of all events of a single category (drug, food, or water intake) that occurred in 10 s bins was taken in a 1000 s time window that surrounded a single drug, food, or water self-administration event.

### Statistical analysis

All of the data was analyzed using Prism 8 software (GraphPad, San Diego, CA, USA). The peristimulus time histogram data were analyzed using two-way repeated-measures analysis of variance (ANOVA) followed by Tukey’s multiple-comparison *post hoc* test. Food intake, water intake, and body weight data were analyzed using two-way ANOVA followed by Tukey’s multiple-comparison *post hoc* test. The data are expressed as mean ± SEM unless otherwise specified.

## Results

### Microstructural analysis of drug, food, and water self-administration events during nicotine self-administration

The microstructural analysis graphs were analyzed as a function of any set of events that occurred 1000 s before and after a single event (i.e., Y *vs*. X). Data were taken during the first day of nicotine access following a 72 h deprivation period after 2 weeks of continuous LgA to maximize the occurrence of nicotine self-administration events^17^ and increase the statistical power of the microstructural meal pattern analysis. Two-way repeated-measures ANOVA was used to compare the effects of time and treatment and time × treatment interactions between animals that self-administered nicotine and animals that self-administered saline.

First, the effect of a single nicotine or saline self-administration event on surrounding nicotine or saline intake events was tested. The ANOVA revealed a significant effect of time on events that surrounded the single self-administration event (*F*_199,1791_ = 1.517, *p* < 0.0001; Fig. 1A), with no significant difference between the number of nicotine or saline events. This lack of significance can be explained by the fact that the animals exhibited a burst of drug-taking events that surrounded a single nicotine self-administration event, thus explaining the significant effect of time; afterward, however, the animals did not lever press as much for nicotine within a short time window as they would for more rewarding drugs, instead taking low doses over a longer period of time.^18^ Next, the effect of nicotine or saline self-administration on surrounding food intake events was tested. Nicotine animals exhibited a significant increase in feeding events that followed a single self-administration event (time × treatment interaction, *F*_199,1791_ = 1.252, *p* = 0.0132; main effect of time, *F*_199,1791_ = 2.033, *p* < 0.0001; Fig. 1B). The *post hoc* analysis showed that nicotine animals exhibited a significant increase in intake from 100 to 340 s following a single nicotine event compared with saline animals. The effect of nicotine or saline self-administration on surrounding water intake events was then tested. Nicotine animals exhibited a significant increase in drinking events following a single self-administration event (time × treatment interaction, *F*_199,1791_ = 1.900, *p* < 0.0001; main effect of time, *F*_199,1791_ = 3.389, *p* < 0.0001; main effect of treatment, *F*_1,9_ = 5.127, *p* = 0.0498; Fig. 1C). The *post hoc* analysis showed that nicotine animals exhibited a significant increase in intake from 10 to 90 s following a single nicotine event compared with saline animals. These results indicate that acute nicotine intake increased feeding and drinking behavior within minutes.

**Figure 1.**
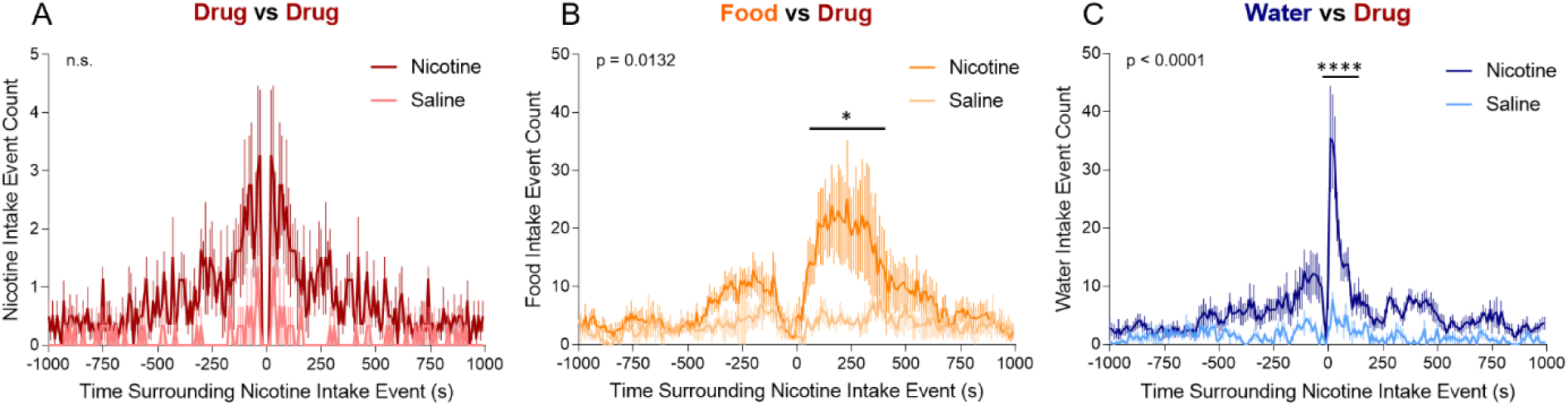
Peristimulus time histograms of drug, food, and water self-administration events that surrounded a single self-administration within a 1000 s time window. (A) Average number of nicotine (dark red) and saline (light red) intake events that surrounded a single nicotine or saline self-administration event, represented by time t = 0. Time × treatment interaction, *p* = 0.8826 (two-way repeated-measures ANOVA). (B) Average number of food intake events that surrounded a single nicotine (dark orange) or saline (light orange) self-administration event. Time × treatment interaction, *p* = 0.0132 (two-way repeated-measures ANOVA). (C) Average number of water intake events that surrounded a single nicotine (dark blue) or saline (light blue) self-administration event. Time × treatment interaction, *p* < 0.0001 (two-way repeated-measures ANOVA). \**p* < 0.05, \*\**p* < 0.01, \*\*\**p* < 0.001, \*\*\*\**p* < 0.0001 (Tukey’s multiple-comparison *post hoc* test).

Next, changes in the duration of each food and drinking bout were analyzed. No significant difference in the duration of a food bout was found between nicotine and saline animals (Fig. 2A). No significant difference in the duration of a water bout was found between nicotine and saline animals (Fig. 2C). However, nicotine animals exhibited a significant decrease in water intake following a single feeding event (time × treatment interaction, *F*_199,1791_ = 1.529, *p* < 0.0001; main effect of time, *F*_159,1791_ = 8.1898, *p* < 0.0001; Fig. 2B). The *post hoc* analysis showed a significantly decrease in water intake from 120 to 290 s following a single food intake event in nicotine animals compared with saline animals.

**Figure 2.**
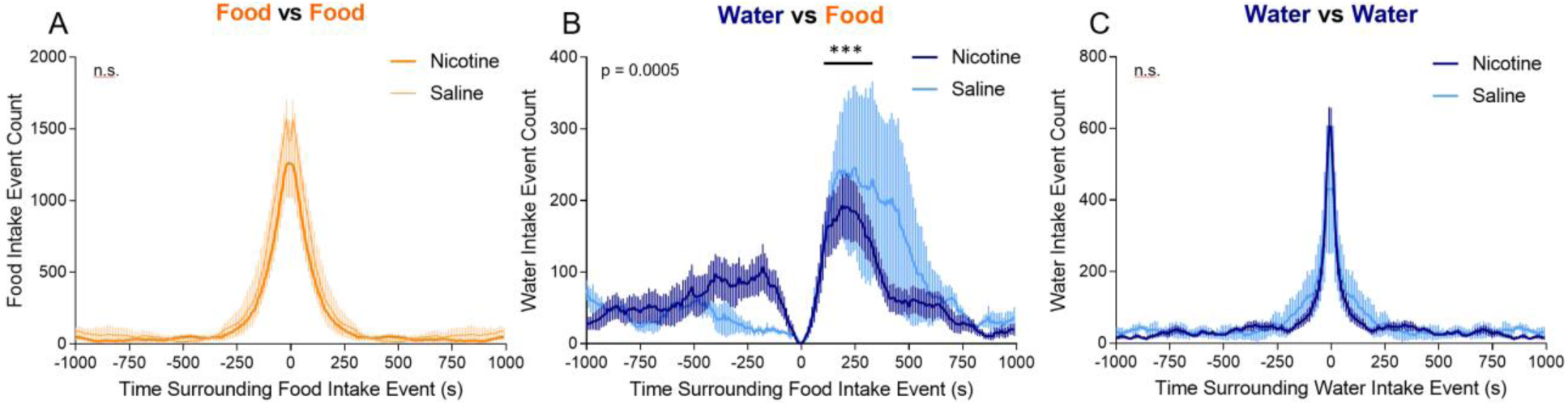
Peristimulus time histograms of food and water intake events that surrounded single food or water intake events within a 1000 s time window. (A) Average number of food intake events that surrounded a single food intake event in rats that self-administered nicotine (dark orange) or saline (light orange). Time × treatment interaction, *p* > 0.9999 (two-way repeated-measures ANOVA). (B) Average number of water intake events that surrounded a single food intake event in rats that self-administered nicotine (dark blue) or saline (light blue). Time × treatment interaction, *p* = 0.0005 (two-way repeated-measures ANOVA). (C) Average number of water intake events that surrounded a single water intake event in rats that self-administered nicotine (dark blue) or saline (light blue). Time = treatment interaction, *p* = 0.9803 (two-way repeated-measures ANOVA). \**p* < 0.05, \*\**p* < 0.01, \*\*\**p* < 0.001, \*\*\*\**p* < 0.0001 (Tukey’s multiple-comparison *post hoc* test).

### Microstructural analysis of drug, food, and water intake events during passive nicotine or saline administration

To test the effect of self-administration behavior on feeding and drinking patterns that surrounded nicotine intake, saline or nicotine was passively administered intravenously using the MedPC program, which required no lever presses by the animals. The animals were given one infusion of 0.9% saline, one infusion of nicotine (0.03 mg/kg/infusion), or three infusions of nicotine (0.03 mg/kg/infusion) and then allowed to nosepoke for food and water. Two-way repeated-measures ANOVA was used to analyze differences between groups.

Fig. 3A shows peristimulus time histograms for passive administration events. Only the three nicotine infusions are shown in the graph because one saline infusion and one nicotine infusion are represented by time t = 0 on the graph. The effect of passive administration events on surrounding food intake events was tested. Animals that were given three nicotine infusions exhibited a significant increase in food intake following the infusions (time × treatment interaction, *F*_198,1188_ = 2.385, *p* < 0.0001). The *post hoc* test showed a significant increase food intake following three nicotine infusions from 180 to 400 s following the passive infusions (Fig. 3B). Next, the effect of passive administration events on surrounding water intake events was tested. A significant increase in water intake was observed in all of the treatment groups following nicotine or saline infusions (main effect of time, *F*_99,1188_ = 3.898, *p* < 0.0001), with no significant difference between groups (Fig. 3C). These results indicate that passive nicotine administration similarly increased food and water self-administration compared with nicotine self-administration.

**Figure 3.**
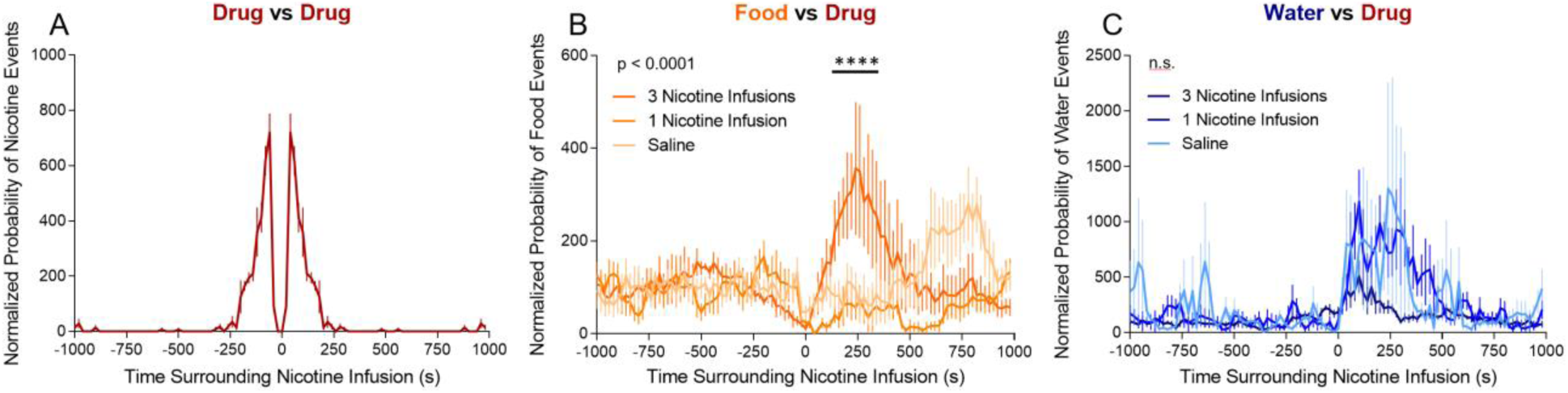
Peristimulus time histograms of drug, food, and water intake events that surrounded passive administration events within a 1000 s time window. (A) Normalized probability of three nicotine infusions. (B) Normalized probability of food intake events in rats following three nicotine infusions (dark orange), one nicotine infusion (orange), or one saline infusion (light orange). Time × treatment interaction, *p* < 0.0001 (two-way repeated-measures ANOVA). (C) Normalized probability of water intake events in rats following three nicotine infusions (dark blue), one nicotine infusion (blue), or 1 saline infusion (light blue). Time × treatment interaction, *p* = 0.7108 (two-way repeated--measures ANOVA). Normalized probability was calculated as the average value for all rats at a single timepoint divided by the average of all events within the 1000 s time window multiplied by 100. \**p* < 0.05, \*\**p* < 0.01, \*\*\**p* < 0.001, \*\*\*\**p* < 0.0001 (Tukey’s multiple-comparison *post hoc* test).

Changes in feeding and drinking bout duration were also analyzed. No significant difference in feeding bouts was observed between treatment groups (Fig. 4A). No significant difference in water intake that surrounded a single feeding event was observed between treatment groups (Fig. 4B). No significant difference in drinking bouts was observed between treatment groups. These results confirmed the feeding and drinking bout data in animals that self-administering nicotine or saline.

**Figure 4.**
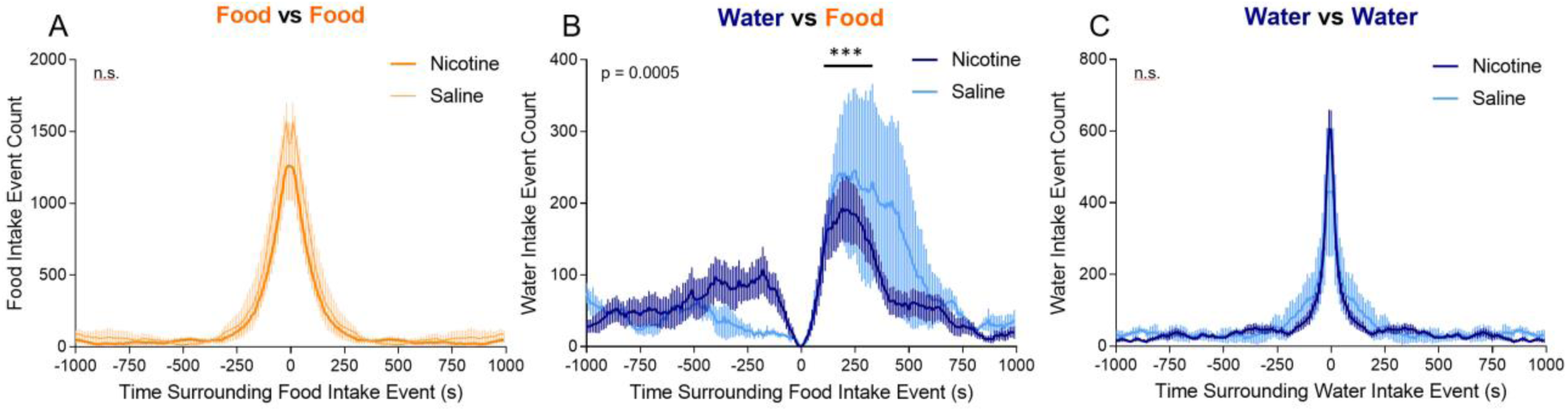
Peristimulus time histograms of food and water intake events that surrounded passive administration events within a 1000 s time window. (A) Normalized probability of food intake events compared with a single food intake event in rats that were given three nicotine infusions (dark orange), one nicotine infusion (orange), or one saline infusion (light orange). Time × treatment interaction, *p* = 0.6658 (two-way repeated-measures ANOVA). (B) Normalized probability of water intake events compared with a single food intake event in rats that were given three nicotine infusions (dark blue), one nicotine infusion (blue), or one saline infusion (light blue). Time × treatment interaction, *p* > 0.9999 (two-way repeated-measures ANOVA). (C) Normalized probability of water intake events compared with a single water intake event in rats that were given three nicotine infusions (dark blue), one nicotine infusion (blue), or one saline infusion (light blue). Time × treatment interaction, *p* > 0.9999 (two-way repeated-measures ANOVA). Normalized probability was calculated as the average for all rats at a single timepoint divided by the average of all events within the 1000 s time window multiplied by 100.

### Macrostructural analysis of feeding behavior during nicotine self-administration

To evaluate the long-term effect of nicotine self-administration on body weight and food and water intake, the rats were given extended access to nicotine self-administration for 6 weeks, during which they received 4 days of 23 h nicotine self-administration (ON) and 3 days without nicotine or saline (OFF) in their home cages with *ad lib* access to food and water. Two-way ANOVA was used to compare the effects of time and treatment and time × treatment interactions between nicotine self-administration animals and saline animals.

The effect of nicotine on body weight was tested. A significant progressive increase in body weight was observed throughout the experiment (main effect of time, *F*_50,300_ = 85.73, *p* < 0.0001). Nicotine self-administration decreased body weight gain compared with saline self-administration (time × treatment interaction, *F*_50,300_ = 2.683, *p* < 0.0001; Fig. 5A). The *post hoc* analysis showed a significant decrease in body weight in nicotine animals compared with saline animals following day 15 of the self-administration paradigm, and this decrease persisted until the end of the experiment. The effect of nicotine on body weight was further examined during a single cycle of the intermittent access paradigm. During the self-administration (ON) phase of the cycle, nicotine animals exhibited a significant decrease in weight gain (time × treatment interaction, *F*_5,30_ = 4.846, *p* = 0.0023; main effect of time, *F*_5,30_ = 7.313, *p* = 0.0001). The *post hoc* analysis showed that nicotine animals exhibited a decrease in body weight gain on days 3 and 4 compared with saline animals However, this decrease in body weight was abolished during the abstinence (OFF) phase of the cycle; by the end of the abstinence phase, body weight was not significantly different between nicotine and saline animals (Fig. 5B).

**Figure 5.**
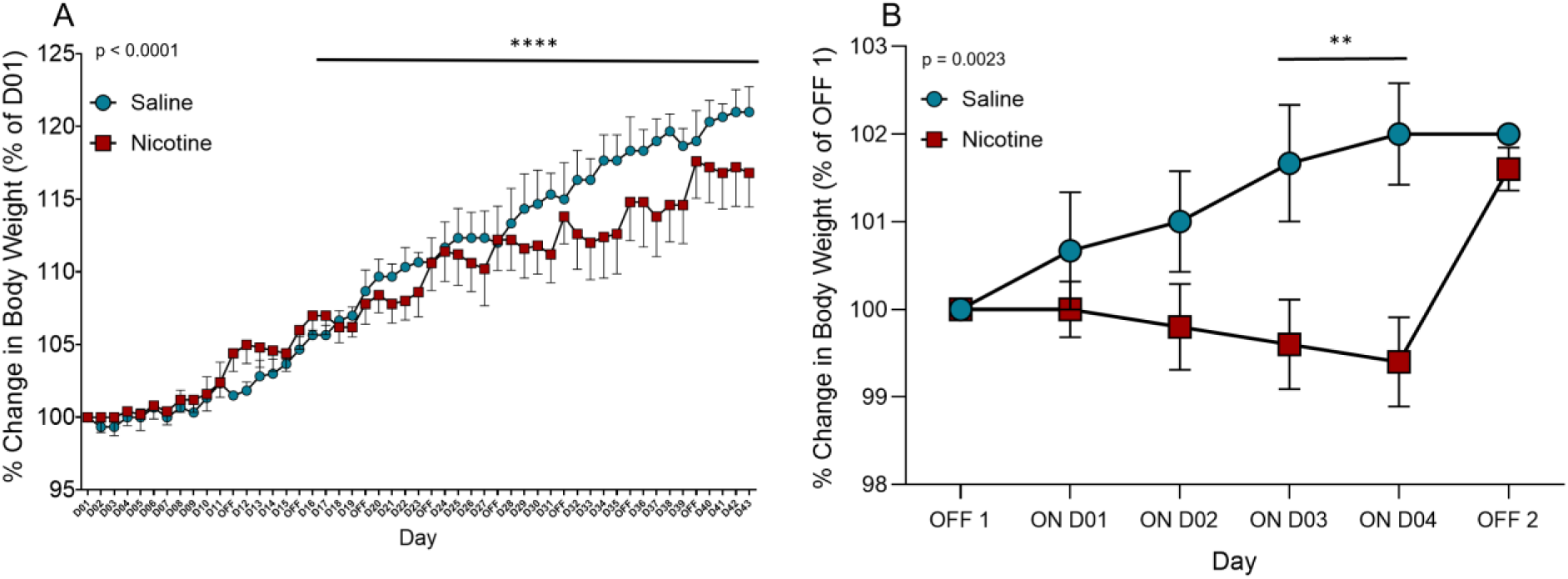
Changes in body weight in nicotine *vs*. saline rats. (A) Average change in body weight in nicotine (red) and saline (blue) rats over the course of the nicotine self-administration paradigm. The data are expressed as a percent change in body weight relative to baseline. Time × treatment interaction, *p* < 0.0001 (two-way repeated-measures ANOVA). (B) Average change in body weight in nicotine (red) and saline (blue) rats during one cycle of the nicotine self-administration paradigm. The data are expressed as a percent change in body weight relative to baseline. Time × treatment interaction, *p* = 0.0023 (two-way repeated-measures ANOVA). \**p* < 0.05, \*\**p* < 0.01, \*\*\**p* < 0.001, \*\*\*\**p* < 0.0001 (Tukey’s multiple-comparison *post hoc* test).

The animals were tested on standard chow and tap water to investigate changes in food and water intake across the self-administration paradigm. Two-way ANOVA was used to compare differences in intake between nicotine and saline animals during the self-administration (ON) and abstinence (OFF) phases. Nicotine animals exhibited a significant increase in total daily chow intake during the abstinence phase compared with saline animals (time × treatment interaction, *F*_1,12_ = 7.255, *p* = 0.0195; main effect of time, *F*_1,12_ = 9.578, *p* = 0.0093; Fig. 6A). During the self-administration phase, nicotine animals exhibited a trend toward a decrease in food intake compared with saline animals (*p* = 0.0731, *t* = 2.170, Student’s *t*-test). No significant difference in total daily water intake during the nicotine ON and OFF phases was observed between nicotine and saline animals (Fig. 6B). These results confirm previous findings that chronic nicotine intake leads to a decrease in body weight gain during nicotine use and an increase in body weight gain and an increase in feeding during nicotine cessation.

**Figure 6.**
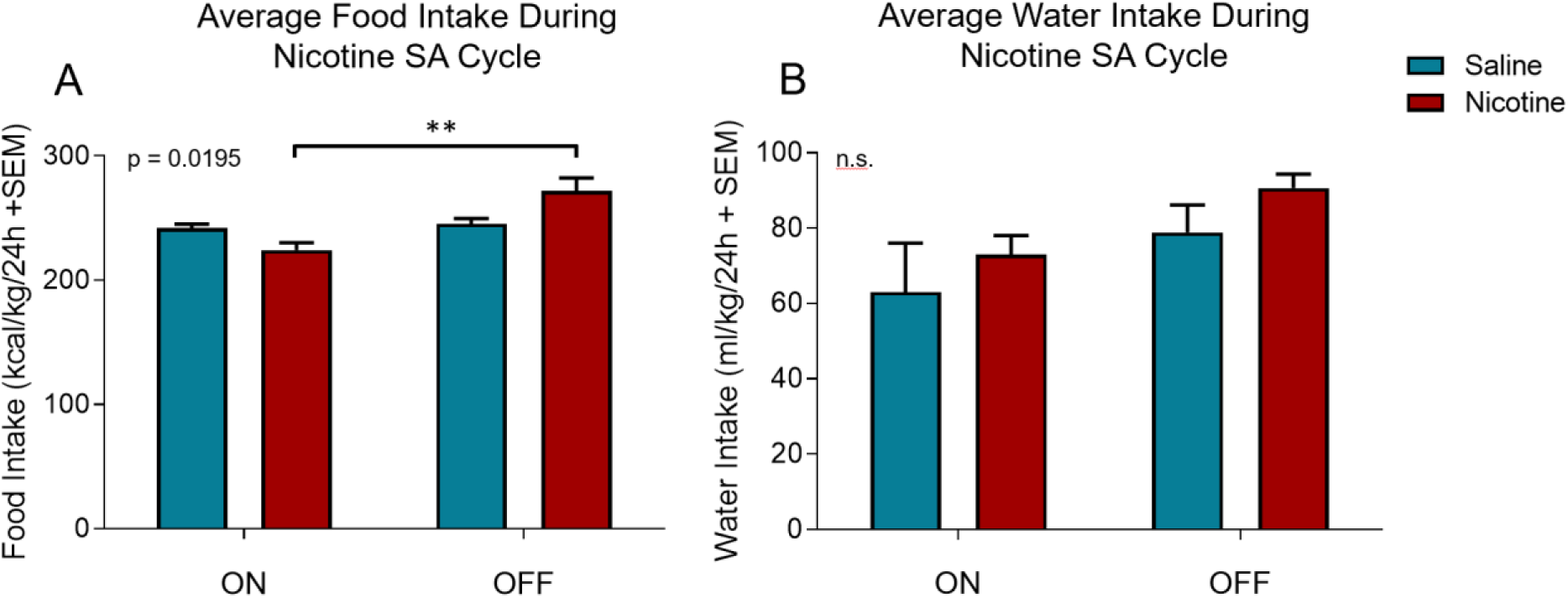
Average food and water intake in nicotine *vs*. saline rats. (A) Average daily food intake in nicotine (red) *vs*. saline (blue) rats during the nicotine ON and OFF phases. The data are expressed as calories consumed normalized to body weight in a 24 h period. Time × treatment interaction, *p* = 0.0195 (two-way ANOVA). (B) Average daily water intake in nicotine (red) *vs*. saline (blue) rats during the nicotine ON and OFF phases. The data are expressed as milliliters of water consumed normalized to body weight in a 24 h period. Time × treatment interaction, *p* = 0.9173 (two-way ANOVA). \**p* < 0.05, \*\**p* < 0.01, \*\*\**p* < 0.001, \*\*\*\**p* < 0.0001 (Tukey’s multiple-comparison *post hoc* test).

## Discussion

Microstructural analyses of behavior during nicotine self-administration showed that rats exhibited a significant increase in food and water intake within minutes of nicotine self-administration compared with saline-self-administration. This increase in feeding behavior was similarly seen in animals following the passive administration of nicotine but not saline. No change in individual feeding or drinking bouts was observed in animals that self-administered nicotine or saline or passively received infusions of nicotine or saline. However, with extended nicotine use, the animals exhibited a decrease in body weight gain and an increase in feeding during nicotine abstinence. These data suggest a paradoxical effect of nicotine on feeding that has not been previously described.

The microstructural analysis of feeding behavior during self-administration showed that the animals exhibited a five-fold increase in food intake that lasted for approximately 4 minutes following a single nicotine intake event. One possibility is that this was a nonspecific effect that was attributable to a general increase in operant behavior. However, this effect was shown to be nicotine-specific, in which the increase in food intake was not observed following saline self-administration. The increase in food intake was also observed following three but not one passive infusions of nicotine, thus demonstrating that the pharmacological effect of nicotine was responsible for the increase in food intake, and that this effect is dose-dependent. These results refute the notion that nicotine is solely an anorectic drug and instead suggest that the initial pharmacological properties of nicotine are anti-anorectic. The present results are consistent with a study by Perkins that showed that smokers actually reported an increase in caloric intake after acute nicotine exposure.^11^

The overall structure of each food and water bout was not affected by nicotine. These results are consistent with previous studies that analyzed meal patterns in rodents,^19,20^ suggesting that nicotine does not affect the bout duration or size but affects the probability that a food or water bout will occur immediately after nicotine administration. Surprisingly, nicotine animals exhibited a 20% decrease in water intake following a food intake event during self-administration. However, this decrease in water intake following a food intake event was not observed in animals following passive nicotine administration. One potential explanation for this discrepancy is that this effect was nonspecific and caused by changes in operant behavior.

The macrostructural analysis data showed that following chronic nicotine use, animals exhibited a 5% decrease in body weight gain compared with saline animals. They also exhibited a 20% increase in food intake during nicotine abstinence compared with when nicotine was available. During each cycle of the intermittent access paradigm, body weight decreased in nicotine animals during the end of the self-administration (ON) phase and robustly increased by the end of the abstinence (OFF) phase. These results support our hypothesis that nicotine intake decreases feeding and weight gain and are consistent with previous studies that reported a decrease in body weight gain during chronic nicotine self-administration and a robust increase following nicotine cessation.^4,5^ The increase in food intake in nicotine animals during abstinence from nicotine is consistent with previous studies in both animals and humans.^21,22^ We found that water intake did not change with chronic nicotine use, which is consistent with previous studies that reported that chronic nicotine did not alter water intake in animals.^23,24^

The opposite effects of acute and chronic nicotine on feeding behavior may be explained by Perkins’ theory that nicotine alters the homeostatic set point of body weight. Perkins and colleagues showed that although body weight decreases with chronic nicotine use in humans, eating increases following acute nicotine intake.^11^ The metabolic effects of nicotine may contribute to a change in body weight set point, with consequent compensatory changes in caloric intake.^25^ The present results are consistent with Perkins’s theory and provide evidence that nicotine interacts with body metabolism to elicit an acute increase in food intake while also leading to a decrease in body weight over time.

In summary, the present study found that nicotine first produced anti-anorectic effects before producing long-term anorexigenic effects. These results challenge the dogma that nicotine is solely an anorexigenic drug and paradoxically suggest that the anorexigenic effects of nicotine may be a long-term consequence of the acute anti-anorectic effects of nicotine. Further studies are needed to understand the cellular and molecular mechanisms of both the anorectic and anti-anorectic effects of nicotine. A better understanding of the acute and long-term effects of nicotine on feeding behavior may contribute to the development of alternative strategies to treat tobacco use disorder and obesity.

## Acknowledgments

The authors thank Michael Alan Arends for proofreading the manuscript.

